# Tonic ERK signaling regulate the basal lactate production in prepubertal rat Sertoli cell

**DOI:** 10.1101/2024.12.18.629288

**Authors:** Mukesh Gautam, Bhola Shankar Pradhan, Subeer S Majumdar

## Abstract

Sertoli cells are the major nurse cells in the testis which regulate the division and differentiation of germ cell by secreting various factors like inhibin B, transferrin and lactate. In this study, we demonstrated that inhibition of extracellular signal-regulated kinase (ERK) leads to an increase in the basal level of lactate. This increase in lactate production was due to the rise in lactate dehydrogenase (LDH) activity and glucose uptake, simultaneously. Moreover, an increase in lactate production wasnot due to the change in mRNA level of *GLU1*, *GLUT3* and *LDHA*. Importantly, the lactate production was independent of the level of cAMP suggesting that the tonic signaling of ERK is important for the growth factor mediated lactate production. These findings suggest that ERK signaling inhibits lactate production in the absence of any stimulant in rat Sertoli cells to maintain metabolic homeostasis.

## Introduction

Sertoli cell (Sc) is the major nurse cell in the testis which supports the division and differentiation of germ cell by secreting various factors. One of their important functions is to convert glucose to lactate, which is the preferred energy source for round spermatids (Hall and Mita 1984; Mita *et al*. 1982). 95.8% glucose uptake by Sc is converted into L-Lactate (Robinson and Fritz 1981). Interestingly, lactate prevents male germ cell apoptosis in human testis (Erkkilä *et al*. 2002). FSH significantly increases lactate production by 16-20 days old Sc (Mita et al. 1982). Lactate production by Sc is primarily regulated by follicle stimulating hormone (FSH), insulin, insulin growth factor I (IGF I) as well as various locally produced paracrine/autocrine factors like transforming growth factor β (TGF β), epidermal growth factor (EGF), basic fibroblast growth factor (bFGF), and interleukin 1α (IL-1α) (Meroni et al. 2019; Pitetti et al. 2013; Galardo et al. 2008; Riera et al. 2007).

LDH is a well characterized isozyme system that is involved in the interconversion of pyruvate and lactate. These isozymes are encoded by three different genes, ldh a (muscle type), ldh b (heart type), and ldh c (testis type) (Li 1989). The latter gives rise to LDH C4 isozyme present only in the mature testis specifically in spermatozoa (Markert et al. 1975). The combinations of the other two gene products result in four tetrameric LDH isozymes, LDH-1 (B4), LDH-2 (A1B3), LDH-3 (A2B2), LDH-4 (A3B1), and LDH-5 (A4), which are present in variable proportions in different somatic tissues including rat Sc (Santiemma et al. 1987).

These factors act through various signal transduction pathways to regulate lactate production in Sc. For example, FSH binding to its receptor (FSHR) leads to activation of adenylate cyclase that results in the consequent increase in cAMP levels and PKA activation which leads to lactate production (Meroni et al. 2002; Walker and Cheng, 2005). IL-1β stimulate lactate production in rat Sc via PI3K/PKB and ERK1/2 pathways (Riera et al. 2007). PI3K activity participates in FSH regulation of Sc cell lactate production (Meroni et al. 2002). MAPK- and PI3K/PKB-dependent pathways participate in bFGF regulation of Sc lactate production (Riera et al. 2003). The bFGF leads to an increase in lactate production by upregulating the glucose transport, LDH activity and GLUT1 and LDH A mRNA levels (Riera et al. 2002). Therefore, all these signal transduction pathways are shared by different growth factors to regulate lactate production in rat Sc.

In the absence of these growth factors, Sc produces little amounts of lactate. This suggests that lactate production is tightly linked to energy metabolism which needs elevated level of NADH. Lactate acts as a mitochondrial messenger to stimulate oxidative phosphorylation and that shifts ATP production to the mitochondria (Cai et al. 2023). It activates the electron transport chain (ETC) without being metabolized (Cai et al. 2023). NADH is produced in the mitochondria. The lactate production is initiated when cells have extra glucose intake and high NADH levels in the presence of stimulants. In this study, we investigated the role of constitutively active signaling pathways in regulating the basal levels of lactate production in Sertoli cells. Our data suggests that ERK signaling acts as a gatekeeper for lactate production in rat Sc in the absence of stimulant.

## Material and Methods

### Animals and reagents

Wistar rats (*Rattus norvegicus*) were obtained from the Small Animal Facility of the National Institute of Immunology (New Delhi, India). All animals were housed and used as per the national guidelines provided by the Committee for the Purpose of Control and Supervision of Experiments on Animals. Protocols for the experiments were approved by the Institutional Animal Ethics Committee and the Committee for the Purpose of Control and Supervision of Experiments on Animals. Ovine (o)FSH, rat FSH (NIAMD rat-FSH-I-6), and anti-cAMP antibodies were obtained from National Hormone and Pituitary Program (NHPP), National Institutes of Health (NIH; Torrance, CA). All other reagents, unless stated otherwise, were procured from Sigma Chemical (St. Louis, MO).

### Isolation of Sc

Testes were dissected from rats of 19 days old (prepubertal). Sc were isolated using a sequential enzymatic digestion that has been previously described by us (Bhattacharya et al. 2012; Bhattacharya et al. 2021). Cultures were continued in DMEM-nutrient mixture F-12 Ham (DMEM-F12 HAM) containing 1% FCS for 24 h in a humidified 5% CO2 incubator at 34°C. Next day, cells were washed with prewarmed medium (DMEM-F12 HAM) and cultured further in serum replacement growth factor medium (GF medium) containing 5 µg/ml sodium selenite, 10 µg/ml insulin, 5 µg/ml transferrin, and 2.5 ng/ml epidermal growth factor. On *day 3* of culture, residual Gc, if any, were removed by hypotonic shock by incubating Sc with 20 mM Tris·HCl (pH 7.4) for 3–5 min at 34°C. Sc were then washed twice to remove dead Gc, and the culture was continued further in GF medium. On *day 4* of culture, Sc were given treatment with various inhibitors and o-FSH for 24h and lactate production was analyzed in the spent media and cells were saved in Trizol for mRNA analysis of LDHA, *GLUT1* and *GLUT3*. In another set of similar experiment, cell were harvested to determine LDH activity. For 2-DOG uptake studies, cells were pretreated with U0126 inhibitor for 24 h. Sertoli cell viability was analyzed with trypan blue exclusion assay.

### Lactate determination

The Sc were treated with different inhibitors (U0126, P38, SQ) and FSH for indicated time in the figure legend. Lactate was measured by a standard method involving conversion of NAD+ to NADH determined as the rate of increase of absorbance at 340 nm. A commercial kit from Sigma-Aldrich with a 5% inter-assay coefficient of variation was used.

### LDH activity measurement

Lactate production was determined as described by us previously (Gautam et al. 2016). After incubation of Sertoli cells in the absence or presence of the different inhibitors (U0126, P38, SQ) and FSH, culture media were saved for lactate determinations and cells were disrupted by ultrasonic irradiation in NaCl (0·9%) and centrifuged (15 800 *g*, 10 min). The supernatant was used to measure total LDH activity. Total LDH activity was determined by a routinely used spectrophotometric method (Randox Laboratories, Crumlin, Antrim, UK).

### Measurement of 2-deoxy-glucose (2-DOG) uptake

Glucose import assay was performed as described previously (Chattopadhyay et al. 2017). Glucose import was studied using the uptake of the labeled non-metabolizable glucose analogue 2-DOG. After treatment with different inhibitors (U0126) and FSH, culture medium was discarded, and cells were washed three times with glucose free DMEM at room temperature. Sertoli cells were then incubated at 34 deg C in 0·5 ml glucose free DMEM containing [2,6-3H]-2-DOG (0·5 μCi/ ml) for 30 min. Unspecific uptake was determined in incubations performed in the presence of a 10 000-fold higher concentration of unlabeled 2-DOG. At the end of the incubation period, dishes were placed on ice and washed extensively with ice-cold PBS until no radioactivity was present in the washings. Cells were then dissolved with 0·5 M sodium hydroxide, 0·4% sodium deoxycholate and counted in a liquid scintillation spectrophotometer. Parallel cultures receiving identical treatments to those performed before the glucose uptake assay were destined for protein determination. Results are expressed on Fmol per μg protein basis.

### Cyclic AMP assay

On *day 4* of culture, media from Sc treated with *1*) GF medium containing 0.1 mM IBMX (vehicle), i.e., Control (C); *2*) vehicle containing o-FSH (25, 50, and 100 ng/ml); *3*) vehicle containing U0126; and *4*) vehicle containing o-FSH and U0126 for 1/2 h were used to determine cAMP by radioimmuno assay (RIA) as reported by us previously (Bhattacharya et al. 2012). Similar experiments were done for 24 h without addition of IBMX to the Sc so that it didnot affect the transcription of gene.

### qRT-PCR

The qRT-PCR was performed as described by us previously (Pradhan et al. 2020, Pradhan et al. 2019; Bhattacharya et al. 2019). Total RNA (1µg) isolated from each treatment group was first reverse transcribed using Reverse Transcription (RT) System (Promega Corp, USA) with AMV reverse transcriptase and oligo (dT)_15_ for the single-strand cDNA synthesis. Subsequent PCR reactions (10µl reaction volume) were carried out using 1µl of the RT reaction as template for checking the expression profile of each gene. For each gene, number of PCR cycles in Biorad PCR machine were standardized to detect an acceptable expression level to confirm the findings.

qRT-PCR amplifications were performed using the RealplexS (Eppendorf) in a total volume of 10 µl (1 or 2 µl of cDNA, depending upon the abundance of the transcripts), 0.5 µM of each primer, and 5 µl of Power SYBR Green Master Mix (Applied Biosystems). Primers for each gene (target gene as well as internal control Ppia) were validated by a standard curve calculated from the CT values of real-time amplification from serial dilutions of the cDNAs. Primers with an efficiency of 1 ± 0.2 were used. The qRT-PCR reaction started with melting of cDNA at 95°C for 15 min followed by 40 amplification cycles (30 s at 95°C, 45 s at 60°C, and 45 s at 65°C). Melting curve analyses for each gene were performed to detect a specific amplification peak for each gene. The expressions of mRNA of the target genes (FSH-R and AR) were evaluated by the efficiency corrected ΔΔCT method. Hormone (FSH or T) and other signaling agents were calculated by relative fold changes using the 2^ΔCT^ method. The means of at least three individual experiments were evaluated for each treatment group for the target gene. Primers used are described in Table 1.

**Table 1.**
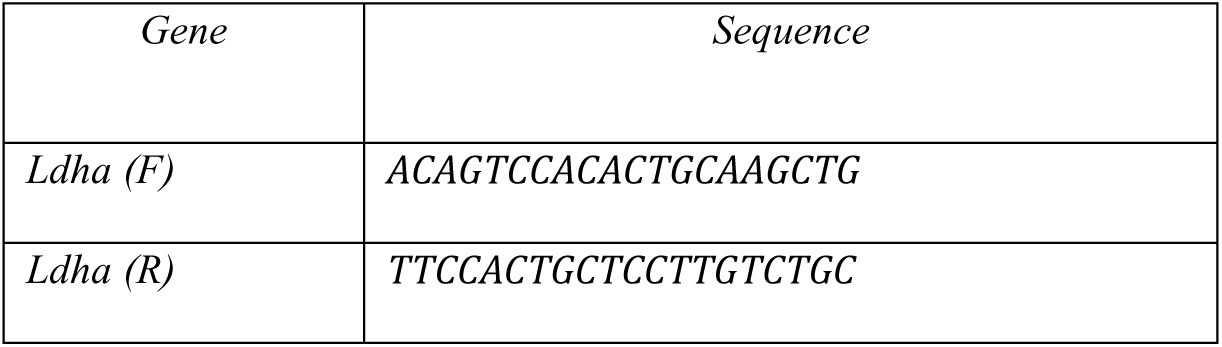
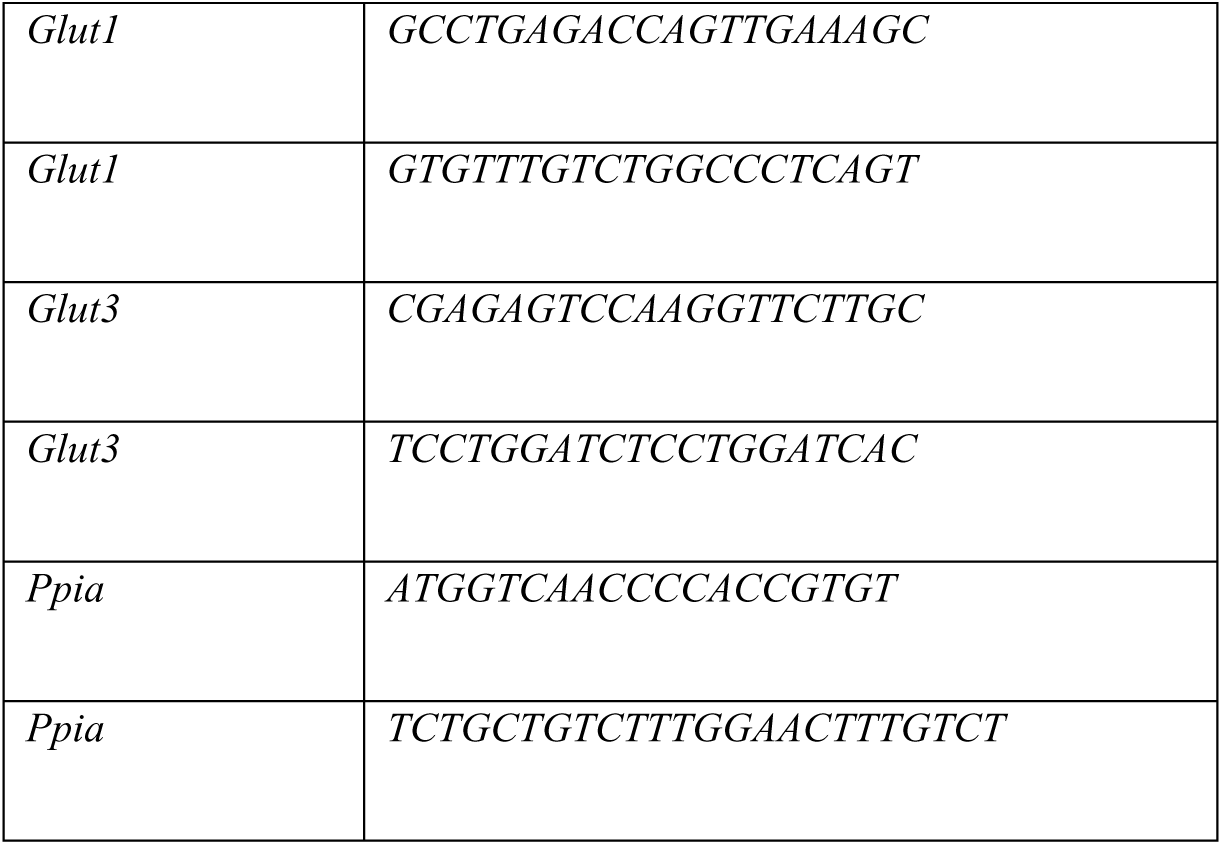
Primers for Q-RT PCR of genes.

### Data representation and statistical analysis

One treatment group comprised three wells within one culture set. At least three such sets of cultures for each age group (performed on different calendar dates) were used to interpret the data. Testes from about 6–10 male rats were pooled for 19-day-old rat Sc cultures. One-way ANOVA followed by Dunnett’s test using the InStat v. 3.0 statistical program (Graphpad Software, San Diego, CA) was used for statistical analyses of the data.

## Results

### ERK inhibition upregulate the basal level of lactate production

To determine the effect of various inhibitors on the production of LDH at basal level, we treated the 19 day old rat Sc with the inhibitor of various pathways. We observed a significant upregulation in basal level of lactate in the media of Sc treated with U0126 as compared to that of control at 24 h (**Fig 1**). This rise in basal level of lactate in the media of U0126 treated Sc was much more than that of FSH treated Sc. Addition of FSH along with U0126 did not alter in the level of lactate production significantly. Similar observations were made with p38 inhibitor (**Suppl Fig 1**). We did not observe any significant rise in the basal level of lactate production with other inhibitor except for ERK inhibitor and p38 inhibitor (**Suppl Fig 1**).

**Figure 1.**
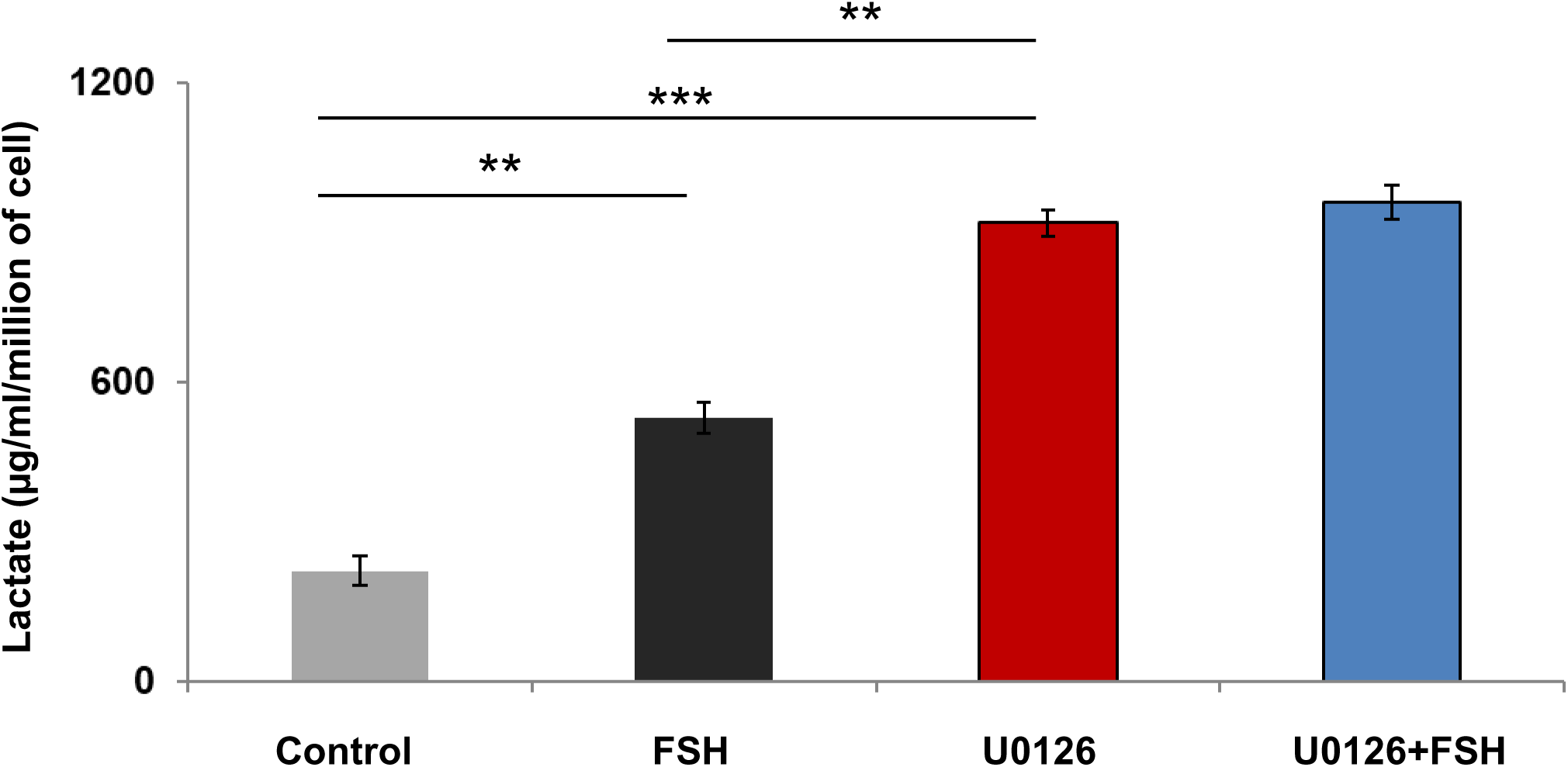
U0126 upregulate the production of lactate. The Sc of 19 days old rat was treated with U0126, FSH and FSH with U0126 for 24h. The media wes collected after the treatment and the lactate were measured. The control cells were treated with vehicles. Each bar represents mean ± SE from ≥3 individual experiments. *P < 0.05, Student t test.

### ERK inhibition increases the LDH activity at basal level

To further investigate whether the increase in the basal level of lactate production due to treatment with U0126 and p38 inhibitor is due to any change in LDH activity, we evaluated the LDH activity in the Sc treated with these inhibitors. We observed a significant rise in the basal level of LDH activity upon treatment with U0126 as compared to that of control at 24 h (**Fig 2**). We did not observe any significant change in the LDH activity in Sc treated with U0126 and FSH in combination as compared to that of U0126 alone. The rise in LDH activity in Sc treated with U0126 was significantly higher than that of control as well as FSH alone. This justified the observation that U0126 treatment alone can increase the lactate production due to the increase in the LDH activity. Another inhibitor of MAPK cascade i.e with P^38^ inhibitor also increased the LDH activity in the Sc suggesting that the increase in LDH activity leads to upregulation in basal level of lactate production (**Suppl Fig 2**).

**Figure 2.**
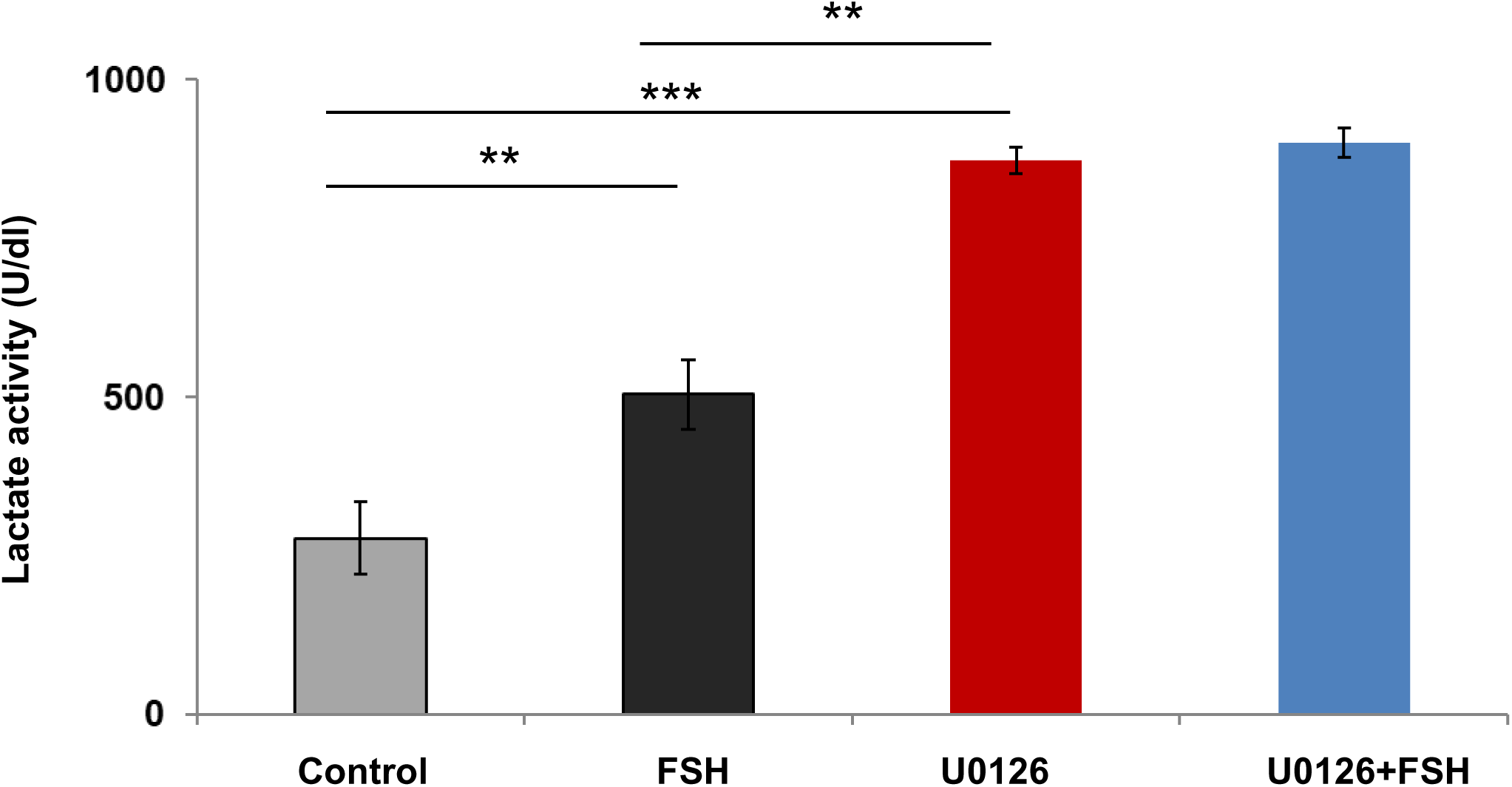
U0126 increases LDH activity. The Sc of 19 days old rat were treated with U0126, FSH and FSH with U0126 for 24h. The control Sc were treated with vehicle only. After treatment, the Sc lysates from different groups were evaluated for LDH activity. Each bar represents mean ± SE from ≥3 individual experiments. *P < 0.05, Student t test.

### ERK inhibition increase the glucose uptake at basal level

To investigate whether the increase in basal level of lactate production (due to treatment with U0126 or p38 inhibitor) affects glucose uptake. The extra glucose enters the glycolytic cycle to produce lactate. We observed a significant rise in the 2-DOG uptake in rat Sc stimulated with U0126 for 24 h (**Fig 3**). The addition of FSH did not have any synergistic effect with the inhibitor in terms of glucose uptake, suggesting that cells glucose uptake may have attained a saturation at 24 h. Moreover, the glucose uptake due to stimulation with U0126 or p38 inhibitor was significantly high as compared to that of FSH alone. These data suggest that the increase in the basal level of lactate production due to ERK inhibitors was due to the upregulation in glucose uptake.

**Figure 3.**
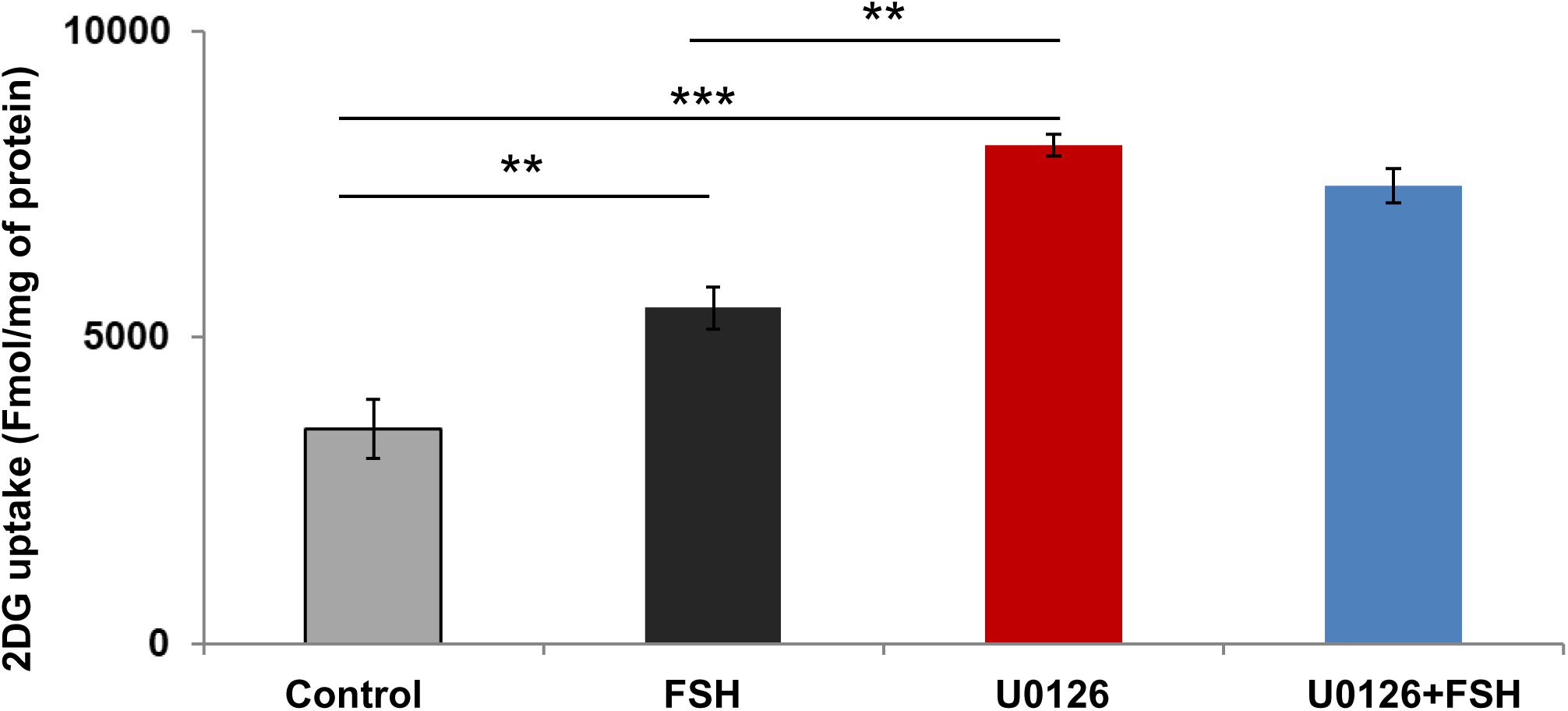
U0126 upregulates the uptake of 2-DOG. The Sc of 19 days old rat were treated with U0126, FSH and FSH with U0126 for 24h. The control Sc were treated with vehicle only. After treatment, Sertoli cells were then incubated at 34 deg C in 0·5 ml glucose free DMEM containing [2,6-3H]-2-DOG (0·5 μCi/ ml) for 30 min and nonspecific uptake was blocked by 10 000-fold higher concentration of unlabeled 2-DOG. After the washing, Sc were counted in a liquid scintillation spectrophotometer. Each bar represents mean ± SE from ≥3 individual experiments. *P < 0.05, Student t test.

### ERK inhibition upregulate the basal level of lactate production without affecting the cAMP level

Since many growth factors like FSH increases the production of cAMP which activate various downstream protein kinases to produce lactate in Sc. Therefore, we measure the level of cAMP as it is a key signaling molecule for lactate production. The treatment of U0126 did not have any significant effect on the level of cAMP as compared to that of control at both ½ h and 24 h (**Fig 4**). Moreover, U0126 significantly upregulated the basal level of lactate production in Sc as compared to that of control as well as FSH alone at both ½ h and 24 h. This upregulation in lactate production in Sc treated with FSH was significantly high at 24 h as compared to that of ½ h. Since FSH leads to an increase in lactate production by increasing the cAMP level which leads to an increase in the transcription of genes related to lactate production such as LDHA, GLUT1 and GLUT3. Our data suggests that at ½ h time point, U0126 leads to an increase in lactate production as compared to that of control or FSH alone suggesting that U0126 is acting directly at the protein level and not on the transcription level for lactate production as ½ h is considered a very small time period for transcription to happen followed by their translation in Sc. This may further suggest that ERK pathway is inhibiting the basal level of lactate production in the absence of any stimulant so that the production of lactate will be regulated.

**Figure 4.**
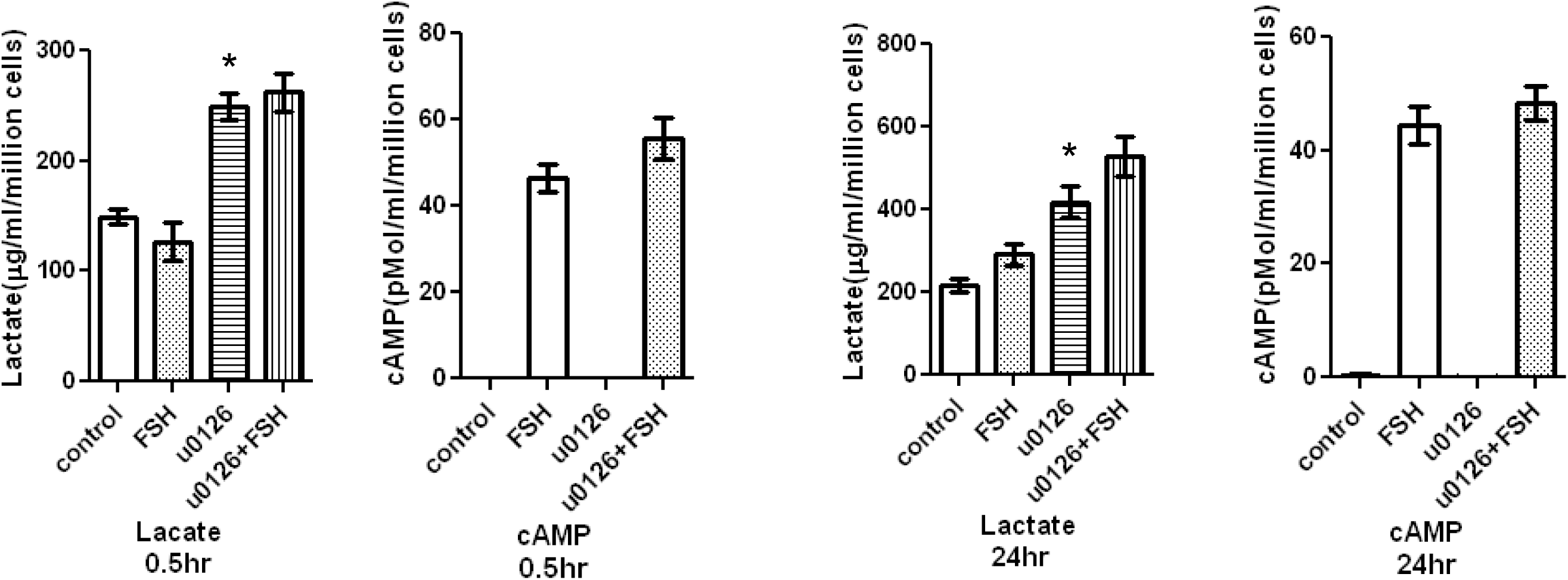
U0126 did not alter the level of cAMP. The Sc of 19 days old rat were treated with U0126, FSH and FSH with U0126 either for ½ h (in presence of IBMX) and for 24 h. The control Sc were treated with vehicle only. After the treatment, cAMP were measured. Simultaneously, lactate level were measured in the corresponding samples. Each bar represents mean ± SE from ≥3 individual experiments. *P < 0.05, Student t test.

### ERK inhibition did not affect the mRNA levels of GLUT1,GLUT3 and LDHA

Since ERK inhibitor did not alter the level of cAMP at both the time point (at ½ h and 24 h), we investigated whether it has any effect on transcriptions of gene related to lactate production such as *LDHA*, *GLUT1* and *GLUT3*. As expected, ERK inhibitor did not alter the mRNA level of *LDHA*, *GLUT1* and *GLUT3* confirming our previous observation that ERK inhibitor is directly affecting at that protein level (**Fig 5**). On the other hand, FSH stimulate the transcription of these genes. We did not observe any significant upregulation in the mRNA level of *LDHA*, *GLUT1* and *GLUT3* in the Sc treated with FSH and U0126 in combination as compared to that of control. This result may explain the lack of synergistic action of FSH and U0126 in terms of lactate production in Sc treated with FSH and U0126 in combination as compared to that of control.

**Figure 5.**
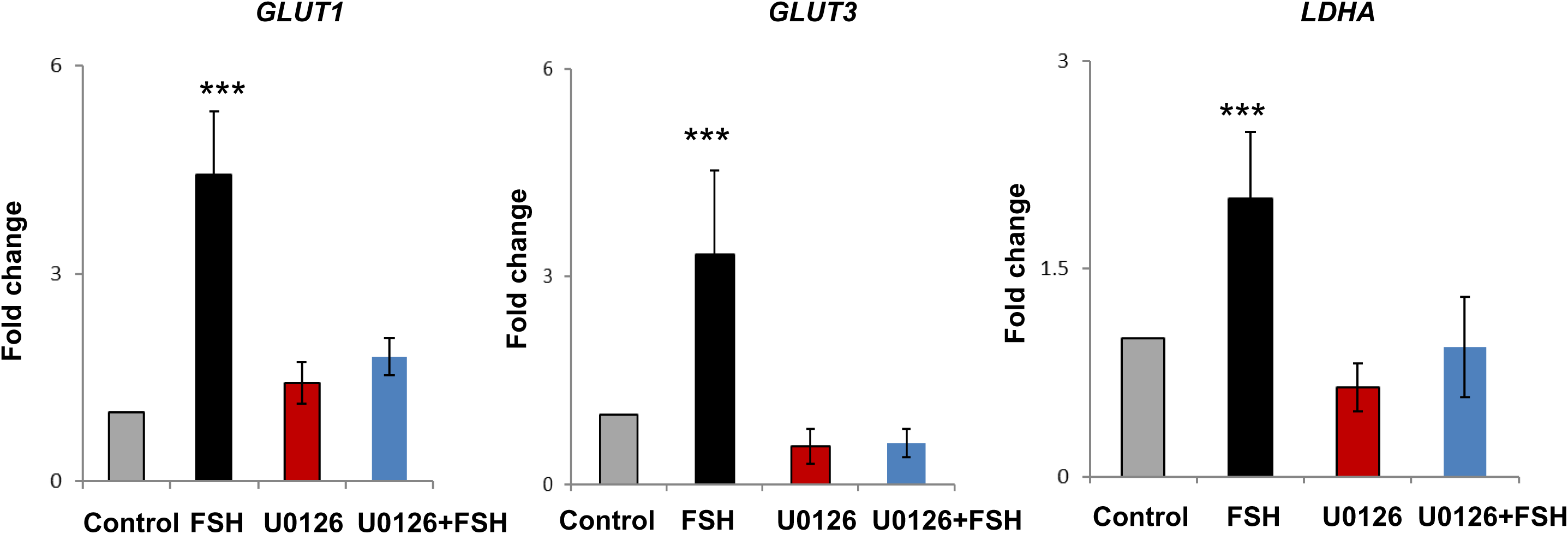
U0126 did not alter mRNA level of *LDHA*, *GLUT1* and *GLUT3.* The Sc of 19 days old rat were treated with U0126, FSH and FSH with U0126 either for ½ h (in presence of IBMX) and for 24 h. The control Sc were treated with vehicle only. After treatment, mRNA expression levels were measured for *LDHA, GLUT1* and *GLUT3.* Each bar represents mean ± SE from ≥3 individual experiments. *P < 0.05, Student t test.

## Discussion

In this study, we demonstrated that ERK signaling acts as a gatekeeper for lactate production in the absence of any stimulant. Our conclusion is based on the observation that two different inhibitors of the MAPK cascade (U0126 and P38 inhibitor) upregulate the production of lactate by stimulating the LDH activity with the simultaneous increase in glucose uptake. This increase in the lactate production was independent of the level of cAMP suggesting that the tonic signaling of ERK is important for the growth factor mediated lactate production.

Lactate is the main source of energy for the spermatids and several signaling molecules regulate its production in Sc. MAPK cascades are arranged as three kinase architecture consisting of a MAPK, a MAPK activator and a MEK activator. In mammalian systems, this MAPK cascades consists of cascades of ERK1/2, JNK and P38. The role of ERK1/2 in regulating various Sc functions have been shown previously (Meroni et al. 2002, Riera et al. 2003). Since lactate production involves high NADH level and high glucose transport into the cells, its production is tightly regulated. Several hormones like FSH, IL1b, IL1α are involved in the regulation of lactate production in Sertoli cells. However, there are no studies about basal levels of lactate production in the absence of stimulant.

In this study, we showed that ERK inhibition upregulated the basal level of lactate production in prepubertal rat Sc and this drastic increase in lactate production was significantly more than that of FSH alone at 24 h time point. We observed a similar result with p38, another MAPK cascade inhibitor suggesting that our observation isnot an artifact. A previous study suggests that ERK1/2 inhibition do not change the basal level of lactate production in rat Sc (Riera et al. 2003). These discrepancies in the lactate production is due to the use of PD 98059 (another ERK inhibitor) and the evaluation time (72h).

The activity of LDH is a function of its conformation and its expression at protein level. Since MAPK pathway increase the basal activity of LDH, we speculate that MAPK pathway (ERK and P^38^ pathway) suppresses the LDH activity in the absence of stimulant to modulate the lactate production. Since glucose uptake is crucial for lactate production as the excess glucose entre the glycolytic pathway, and finally it is converted into lactate. This indicates that ERK inhibition alter the LDH activity and glucose uptake which leads to an upregulation in the basal level of lactate. The increase in glucose uptake without any increase in the mRNA level of *GLUT1* and *GLUT3* is supported by the previous results where short term stimulation of b FGF increases the glucose uptake without any increase in glucose transporter (Riera *et al. 2002)*. Thus, our data suggest that the basal activity of ERK signaling is preventing the lactate production in the absence of stimulant. Interestingly, ERK is dramatically down regulated in 19 day old rat Sc (support the division and differentiation of germ cell) as compared to that of 5 day old rat Sc upon FSH treatment (did not support the division and differentiation of germ cell) (Bhattacharya et al. 2012). This is associated with a significant upregulation in the lactate production in 19 day old rat Sc as compared to that of 5 day old rat Sc upon FSH treatment (Crépieux *et al*. 2001). Thus, our findings may explain that ERK inhibition by FSH leads to an increase in the lactate production in the 19 day old rat Sc.

In summary, we report here the tonic signaling or the constitutively active signaling of ERK modulating the basal level of lactate in 19 day old rat Sc. This finding might have important implications in understanding the pathway regulating lactate production in different cell systems such as skeletal muscle or astrocytes.

## Acknowledgments

We are thankful to all the staff of the Small Animal Facility. Thanks are due to Ram Singh, Dharamvir Singh and Birendar Roy for the technical assistance. We are grateful to the Director, NII for valuable support. We are grateful to Department of Biotechnology, Govt. of India, for providing the financial assistance under grants BT/PR11313/AAQ/01/376/2008, BT/HRD/35/01/01/2010 (TATA Innovation Award) and BT/PR10805/AAQ/1/576/2013. BSP was supported the Polish National Science Centre grant (2020/39/D/NZ5/02004).

## Author Contributions

Experiments were conceived and designed by M.G., B.S.P., and S.S.M. Experiments were performed by M.G. and B.SP. Data of the manuscript was analyzed by M.G. and B.SP. Manuscript was written and reviewed by B.S.P. All authors have read and agreed to the published version of the manuscript.

## References

1. Bhattacharya I, Pradhan BS, Sarda K, Gautam M, Basu S, Majumdar SS. A switch in Sertoli cell responsiveness to FSH may be responsible for robust onset of germ cell differentiation during prepubartal testicular maturation in rats. Am J Physiol Endocrinol Metab. 2012 Oct 1;303(7):E886–98. doi: 10.1152/ajpendo.00293.2012. Epub 2012 Jul 31. PMID: 22850685.

2. Bhattacharya I, Basu S, Pradhan BS, Sarkar H, Nagarajan P, Majumdar SS. Testosterone augments FSH signaling by upregulating the expression and activity of FSH-Receptor in Pubertal Primate Sertoli cells. Mol Cell Endocrinol. 2019 Feb 15;482:70–80. doi: 10.1016/j.mce.2018.12.012. Epub 2018 Dec 21. PMID: 30579957.

3. Bhattacharya I, Sharma SS, Sarkar H, Gupta A, Pradhan BS, Majumdar SS. FSH mediated cAMP signalling upregulates the expression of Gα subunits in pubertal rat Sertoli cells. Biochem Biophys Res Commun. 2021 Sep 10;569:100–105. doi: 10.1016/j.bbrc.2021.06.094. Epub 2021 Jul 5. PMID: 34237428.

4. Cai X, Ng CP, Jones O, Fung TS, Ryu KW, Li D, Thompson CB. Lactate activates the mitochondrial electron transport chain independently of its metabolism. Mol Cell. 2023 Nov 2;83(21):3904–3920.e7. doi: 10.1016/j.molcel.2023.09.034. Epub 2023 Oct 24. PMID: 37879334; PMCID: PMC10752619.

5. Chattopadhyay T, Singh RR, Gupta S, Surolia A. Bone morphogenetic protein-7 (BMP-7) augments insulin sensitivity in mice with type II diabetes mellitus by potentiating PI3K/AKT pathway. Biofactors. 2017 Mar;43(2):195–209. doi: 10.1002/biof.1334. Epub 2017 Feb 10. PMID: 28186649.

6. Crépieux P, Marion S, Martinat N, Fafeur V, Vern YL, Kerboeuf D, Guillou F, Reiter E. The ERK-dependent signalling is stage-specifically modulated by FSH, during primary Sertoli cell maturation. Oncogene. 2001 Aug 2;20(34):4696–709. doi: 10.1038/sj.onc.1204632. PMID: 11498792.

7. Erkkilä K, Aito H, Aalto K, Pentikäinen V, Dunkel L. Lactate inhibits germ cell apoptosis in the human testis. Mol Hum Reprod. 2002 Feb;8(2):109–17. doi: 10.1093/molehr/8.2.109. PMID: 11818513

8. Gautam M, Bhattacharya I, Devi YS, Arya SP, Majumdar SS. Hormone responsiveness of cultured Sertoli cells obtained from adult rats after their rapid isolation under less harsh conditions. Andrology. 2016 May;4(3):509–19. doi: 10.1111/andr.12161. Epub 2016 Mar 18. PMID: 26991307.

9. Galardo MN, Riera MF, Pellizzari EH, Chemes HE, Venara MC, Cigorraga SB, Meroni SB. Regulation of expression of Sertoli cell glucose transporters 1 and 3 by FSH, IL1 beta, and bFGF at two different time-points in pubertal development. Cell Tissue Res. 2008 Nov;334(2):295-304. doi: 10.1007/s00441-008-0656-y. Epub 2008 Sep 19. PMID: 18802725.

10. Hall PF, Mita M. Influence of follicle-stimulating hormone on glucose transport by cultured Sertoli cells. Biol Reprod. 1984 Dec;31(5):863–9. doi: 10.1095/biolreprod31.5.863. PMID: 6097313.

11. Markert CL, Shaklee JB, Whitt GS. Evolution of a gene. Multiple genes for LDH isozymes provide a model of the evolution of gene structure, function and regulation. Science. 1975 Jul 11;189(4197):102-14. doi: 10.1126/science.1138367. PMID: 1138367.

12. Meroni SB, Galardo MN, Rindone G, Gorga A, Riera MF, Cigorraga SB. Molecular Mechanisms and Signaling Pathways Involved in Sertoli Cell Proliferation. Front Endocrinol (Lausanne). 2019 Apr 16;10:224. doi: 10.3389/fendo.2019.00224. PMID: 31040821; PMCID: PMC6476933.

13. Meroni SB, Riera MF, Pellizzari EH, Cigorraga SB. Regulation of rat Sertoli cell function by FSH: possible role of phosphatidylinositol 3-kinase/protein kinase B pathway. J Endocrinol. 2002 Aug;174(2):195–204. doi: 10.1677/joe.0.1740195. PMID: 12176658.

14. Mita M, Hall PF. Metabolism of round spermatids from rats: lactate as the preferred substrate. Biol Reprod. 1982 Apr;26(3):445–55. doi: 10.1095/biolreprod26.3.445. PMID: 7082719.

15. Li SS. Lactate dehydrogenase isoenzymes A (muscle), B (heart) and C (testis) of mammals and the genes coding for these enzymes. Biochem Soc Trans. 1989 Apr;17(2):304–7. doi: 10.1042/bst0170304. PMID: 2753209.

16. Pitetti JL, Calvel P, Zimmermann C, Conne B, Papaioannou MD, Aubry F, Cederroth CR, Urner F, Fumel B, Crausaz M, Docquier M, Herrera PL, Pralong F, Germond M, Guillou F, Jégou B, Nef S. An essential role for insulin and IGF1 receptors in regulating sertoli cell proliferation, testis size, and FSH action in mice. Mol Endocrinol. 2013 May;27(5):814–27. doi: 10.1210/me.2012-1258. Epub 2013 Mar 21. PMID: 23518924; PMCID: PMC5416760.

17. Pradhan BS, Bhattacharya I, Sarkar R, Majumdar SS. Pubertal down-regulation of Tetraspanin 8 in testicular Sertoli cells is crucial for male fertility. Mol Hum Reprod. 2020 Oct 1;26(10):760–772. doi: 10.1093/molehr/gaaa055. PMID: 32687199.

18. Pradhan BS, Bhattacharya I, Sarkar R, Majumdar SS. Downregulation of Sostdc1 in Testicular Sertoli Cells is Prerequisite for Onset of Robust Spermatogenesis at Puberty. Sci Rep. 2019 Aug 7;9(1):11458. doi: 10.1038/s41598-019-47930-x. PMID: 31391487; PMCID: PMC6686024.

19. Riera MF, Galardo MN, Pellizzari EH, Meroni SB, Cigorraga SB. Participation of phosphatidyl inositol 3-kinase/protein kinase B and ERK1/2 pathways in interleukin-1beta stimulation of lactate production in Sertoli cells. Reproduction. 2007 Apr;133(4):763–73. doi: 10.1530/rep.1.01091. PMID: 17504920.

20. Riera MF, Meroni SB, Pellizzari EH, Cigorraga SB. Assessment of the roles of mitogen-activated protein kinase and phosphatidyl inositol 3-kinase/protein kinase B pathways in the basic fibroblast growth factor regulation of Sertoli cell function. J Mol Endocrinol. 2003 Oct;31(2):279–89. doi: 10.1677/jme.0.0310279. PMID: 14519096.

21. Riera MF, Meroni SB, Schteingart HF, Pellizzari EH, Cigorraga SB. Regulation of lactate production and glucose transport as well as of glucose transporter 1 and lactate dehydrogenase A mRNA levels by basic fibroblast growth factor in rat Sertoli cells. J Endocrinol. 2002 May;173(2):335–43. doi: 10.1677/joe.0.1730335. PMID: 12010641.

22. Robinson R, Fritz IB. Metabolism of glucose by Sertoli cells in culture. Biol Reprod. 1981 Jun;24(5):1032–41. doi: 10.1095/biolreprod24.5.1032. PMID: 6268203.

23. Santiemma V, Salfi V, Casasanta N, Fabbrini A. Lactate dehydrogenase and malate dehydrogenase of Sertoli cells in rats. Arch Androl. 1987;19(1):59–64. doi: 10.3109/01485018708986800. PMID: 3122678.

24. Walker WH, Cheng J. FSH and testosterone signaling in Sertoli cells. Reproduction. 2005 Jul;130(1):15–28. doi: 10.1530/rep.1.00358. PMID: 15985628.

