## Supplementary file for "Tonic ERK signaling regulate the basal lactate production in prepubertal rat Sertoli cell"

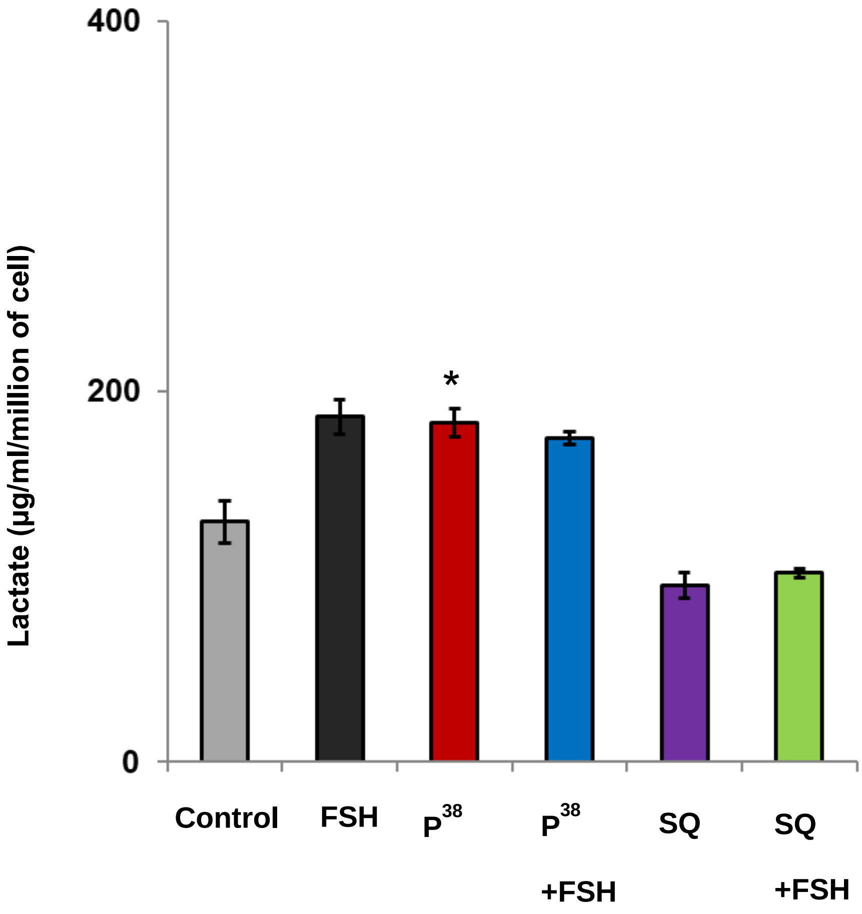


**Supplementary Figure 1. Effect of different inhibitors on the production of lactate**.

The Sc of 19 days old rat were treated with p38, FSH, FSH with p38, SQ and SQ with FSH with p38 for 24h. The media were collected after the treatment and the lactate were measured. The control cells were treated with vehicles. Each bar represents mean ± SE from ≥3 individual experiments. *P < 0.05, one-way ANOVA followed by Dunnett's posttest.

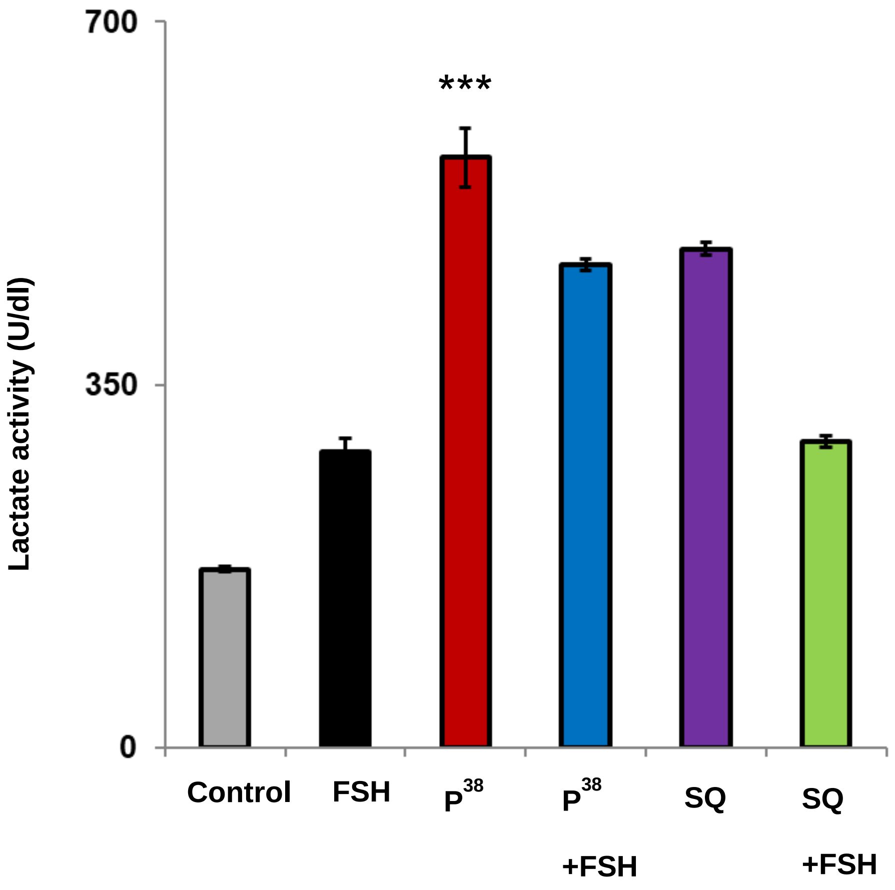


**Supplementary Figure 2. Effect of different inhibitors on the LDH activity**. The Sc of 19 days old rat were treated with p38, FSH, FSH with p38, SQ and SQ with FSH with p38 for 24h. After treatment, the Sc lysates from different groups were evaluated for LDH activity. Each bar represents mean ± SE from ≥3 individual experiments. *P < 0.05, one-way ANOVA followed by Dunnett's posttest.
